# Distinguishing multiple-merger from Kingman coalescence using two-site frequency spectra

**DOI:** 10.1101/461517

**Authors:** Eliot F. Fenton, Daniel P. Rice, John Novembre, Michael M. Desai

## Abstract

Demographic inference methods in population genetics typically assume that the ancestry of a sample can be modeled by the Kingman coalescent. A defining feature of this stochastic process is that it generates genealogies that are binary trees: no more than two ancestral lineages may coalesce at the same time. However, this assumption breaks down under several scenarios. For example, pervasive natural selection and extreme variation in offspring number can both generate genealogies with “multiple-merger” events in which more than two lineages coalesce instantaneously. Therefore, detecting multiple mergers (and other violations of the Kingman assumptions) is important both for understanding which forces have shaped the diversity of a population and for avoiding fitting misspecified models to data. Current methods to detect multiple mergers in genomic data rely primarily on the site frequency spectrum (SFS). However, the signatures of multiple mergers in the SFS are also consistent with a Kingman coalescent with a time-varying population size. Here, we present a new statistical test for determining whether the Kingman coalescent with any population size history is consistent with population data. Our approach is based on information contained in the two-site joint frequency spectrum (2-SFS) for pairs of linked sites, which has a different dependence on the topologies of genealogies than the SFS. Our statistical test is global in the sense that it can detect when the genome-wide genetic diversity is inconsistent with the Kingman model, rather than detecting outlier regions, as in selection scan methods. We validate this test using simulations, and then apply it to demonstrate that genomic diversity data from *Drosophila melanogaster* is inconsistent with the Kingman coalescent.

## Introduction

The genetic diversity within a population reflects its demographic and evolutionary history. Learning about this history from contemporary sequence data is the domain of modern population genetics (see Hahn (2018)). The fundamental tools of the trade are simplified mathematical models, which connect unobserved quantities such as the population size to observable features of genetic data. However, populations are complicated and, moreover, vary in their complications. No simple model can capture the processes governing every species’ evolution, and a misspecified model will generate misleading inferences. It is therefore crucial to understand the limits of population genetics models and to assess when a model is appropriate for a particular data set.

One of the most widely used models is the Kingman coalescent (Kingman 1982a,b; Hudson 1983; Tajima 1983). The Kingman coalescent is a stochastic process that generates gene genealogies: trees representing the patterns of shared ancestry of sampled individuals. Inference methods use these genealogies as latent variables linking demographic parameters to genetic data (Rosenberg and Nordborg 2002). The Kingman coalescent has a number of convenient properties that facilitate both analytical calculations (e.g., Tajima (1989)) and efficient stochastic simulations (e.g., Hudson (2002)): tree topologies are independent of waiting times; waiting times are generated by a Markov process; and neutral mutations are modeled as a Poisson process conditionally independent of the tree. Moreover, the model can be extended to study a variety of biological phenomena including recombination, population structure, and variation in sex ratios or ploidy (see generally Wakeley (2009)).

An important application of the Kingman coalescent is inferring historical population sizes from genetic data (Schraiber and Akey 2015). In its simplest form, the model has a single parameter, the coalescent rate, which determines the branch lengths of genealogies (Kingman 1982a). Under many conditions, the coalescent rate is inversely proportional to the population size (Kingman 1982b). Accordingly, a growing or shrinking population may be modeled by a time-varying coalescence rate (Griffiths and Tavaré 1994, 1998). Patterns of genetic diversity depend on the ratio of the coalescent rate to other evolutionary rate parameters. For example, the *site frequency spectrum* (SFS)— the number of mutations segregating at different frequencies in a sample—is determined by the ratio of the mutation rate to the (time-varying) coalescent rate. Kingman-coalescent–based inference methods solve the inverse problem of determining the population size history that best explains particular features of the data, such as the SFS (e.g., Bhaskar *et al*. (2015)) or variations in heterozygosity along a chromosome (e.g., Li and Durbin (2011)).

A serious problem for this class of inference methods is that different models of evolution generate different relationships between historical population sizes and genetic diversity. For example, one of the basic assumptions of the Kingman coalescent is that natural selection is negligible in determining the distribution of genealogies. When this assumption is violated, Kingman-based inference methods are misspecified (Gillespie 2000a,b, 2001). For instance, when a beneficial mutation increases rapidly in frequency, it distorts the genealogies at nearby sites (see e.g., Coop and Ralph (2012)). If these “selective sweeps” occur regularly, they may be the dominant factor determining the distribution of genealogies. In this case, the average coalescent rate is proportional to the number of beneficial mutations introduced per generation, which is itself *directly*, rather than inversely, proportional to the population size. It follows that the relationship between the population size and the expected number of neutral mutations in a sample is inverted: larger populations will be less diverse than smaller populations.

While the example above is extreme, it is well established that violations of the neutrality assumption can distort or mask the signatures of population size changes. For example, Schrider *et al*. (2016) and Johri *et al*. (2021) demonstrated that several popular inference methods give misleading results in the presence of selective sweeps and background selection. In a similar vein, Cvijović *et al*. (2018) showed that reduction of genetic diversity by purifying selection is accompanied by distortions in the SFS, leading to a false signal of population growth. Moreover, genomic evidence from multiple species suggests that such violations of neutrality may be widespread (Sella *et al*. 2009; Corbett-Detig *et al*. 2015; Kern and Hahn 2018; Johri *et al*. 2020).

An important extension of the Kingman coalescent is a family of models known as *multiple-merger coalescents* (Pitman 1999; Sagitov 1999; Donnelly and Kurtz 1999; Eldon 2016), which arise in a variety of contexts both with and without selection. Whereas in the Kingman coalescent lineages may coalesce only pairwise, multiple-merger coalescents permit more than two lineages to coalesce in a single event. The more general class of simultaneous-multiple-merger coalescents (Schweinsberg 2000; Möhle and Sagitov 2001; Sagitov 2003) permits more than one distinct multiple-merger event at the same time. Multiple-merger and simultaneous-multiple-merger models are relevant for species with “sweepstakes” reproductive events (Eldon and Wakeley 2006; Sargsyan and Wakeley 2008), fat-tailed offspring number distributions (Schweinsberg 2003; Hallatschek 2018), recurring selective sweeps at linked sites (Durrett and Schweinsberg 2005; Coop and Ralph 2012), rapid adaptation (Neher and Hallatschek 2013; Desai *et al*. 2013), and purifying selection at sufficiently many sites (Seger *et al*. 2010; Nicolaisen and Desai 2012; Good *et al*. 2014).

In each of these contexts, the coalescent timescale is not necessarily proportional to the population size. For example, with fat-tailed offspring distributions, the rate of coalescence is a power law in the population size (Schweinsberg 2003), while with linked sweeps it is determined by the rate of linked sweeps, as described above (Durrett and Schweinsberg 2005). In these settings, interpreting the level of genetic diversity in terms of an “effective population size” is misleading, and inferences based on the Kingman coalescent may be qualitatively and quantitatively incorrect.

It is therefore important to determine whether the Kingman model is appropriate for a given data set before performing demographic inference. This task is distinct from “selection scan” methods designed to detect particular regions of the genome that are under selection (see Vitti *et al*. (2013)). Selection scan methods typically assume that most of the genome is evolving neutrally and that the genome-wide distribution of summary statistics reflects demographic factors. Genomic regions that are outliers from this distribution are presumed to be under selection. In contrast, we are interested in detecting when the genome-wide background *itself* is not well-modeled by the Kingman coalescent.

There has been much recent interest in identifying in genomic data departures from the Kingman coalescent caused by multiple mergers. One approach is to use the SFS as a summary statistic. To this end, Birkner *et al*. (2013), Blath *et al*. (2016), and Spence *et al*. (2016) derived methods for computing the expected SFS of (simultaneous) multiple-merger coalescents. Further, Eldon *et al*. (2015) showed that it is possible to use the SFS to distinguish beta and Dirac (multiple-merger) coalescents from Kingman coalescents with strictly exponential or algebraic growth. Koskela (2018) and Koskela and Wilke Berenguer (2019) extended this work and used the SFS to distinguish multiple mergers caused by selection from those caused by sweepstakes reproduction. In a related approach, Rödelsperger *et al*. (2014) detected widespread linked selection in the nematode *Pristionchus pacificus* by demonstrating that the SFS is non-monotonic, a signature of multiple mergers (Neher and Hallatschek 2013; Birkner *et al*. 2013). More recently, Freund and Siri-Jégousse (2021) have introduced a new statistic, the minimum observable clade size, and used it, along with SFS-derived statistics, to discriminate between several coalescent models (including multiple-merger and Kingman coalescents both with and without population growth) using an approximate Bayesian computation (ABC) framework. Several other recent papers have used related combinations of linkage disequilibrium and SFS-derived statistics in a similar ABC framework to detect evidence for multiple-merger genealogies (Menardo *et al*. 2021) or to jointly infer the action of demography and selection (Johri *et al*. 2020, 2021; Lepers *et al*. 2021).

However, methods that derive their power primarily from the SFS are limited in their ability to distinguish multiple mergers from general models of population-size change. While previous work has demonstrated that the SFS does contain information that can discriminate multiple-mergers from particular forms of the Kingman coalescent, a Kingman coalescent with a more general model of population size change can accurately fit many aspects of the multiple-merger SFS (Myers *et al*. 2008; Bhaskar and Song 2014). This fundamentally limits the ability to discriminate between population models using SFS-based statistics alone. The non-monotonic SFS identified by Rödelsperger *et al*. (2014) is a more robust signature of multiple mergers, but identifying that the SFS increases at high frequencies requires both knowledge of the ancestral allele at each site and a large enough sample size to accurately sample rare, high-frequency alleles, and either condition may be violated in real-world data.

Here, we propose that statistics based on the two-site frequency spectrum (2-SFS)—the generalization of the SFS to pairs of nearby sites (Hudson 2001; Ferretti *et al*. 2018)—are useful for distinguishing between the Kingman coalescent with population growth and multiple-merger coalescents. This is fundamentally different from approaches based primarily on the single-site SFS (e.g. Birkner *et al*. (2013); Blath *et al*. (2016); Spence *et al*. (2016); Eldon *et al*. (2015); Freund and Siri-Jégousse (2021)) because 2-SFS-based statistics depend on tree topologies and coalescent rates in a manner unique from SFS-based statistics. Thus these 2-SFS statistics introduce new information not contained in the SFS that can be used to discriminate models that produce same SFS. Furthermore, these statistics may be calculated efficiently from single-nucleotide-polymorphism (SNP) data, do not require recombination maps or ancestral allele identification, and are informative even with small sample sizes. Together, these properties make the 2-SFS useful for demographic model-checking in a wide range of species.

In this paper, we show that 2-SFS-based statistics can be used to discriminate Kingman from non-Kingman coalescence. By validating with simulations, we demonstrate high power to reject incorrect Kingman population-size-change models for biologically realistic sample sizes. We present a `Snakemake` pipeline for analyzing real-world population data and demonstrate our pipeline using genomic data from *Drosophila melanogaster* (Lack *et al*. 2015).

## Definitions and Background

Following the notation of Fu (1995), we define the SFS of a sample of *n* haploid genomes as *ξ*, where *ξ*_*i*_ is the fraction of sites containing a mutation with derived allele count *i* in the sample (1 ≤ *i* ≤ *n* − 1). When the ancestral allele is unknown, mutations at frequency *n* − *i* are indistinguishable from mutations at frequency *i*, and the folded SFS, *η*, is used instead, where *η*_*i*_ is the fraction of sites with minor allele count *i* in the sample, { *η*_*i*_ = *ξ*_*i*_ + (1 − *δ*_*i,n*−*i*_) *ξ*_*n*−*i*_ : 1 ≤ *i* ≤ ⌊*n*/2⌋}. The SFS and folded SFS can be calculated from a set of SNPs without knowing the physical location of the SNPs.

In contrast, the 2-SFS, *ϕ*, is a statistic of *pairs* of sites. We define the 2-SFS, {*ϕ*_*ij*_ (*d*) : *d >* 0; 1 ≤ {*i, j*} ≤ *n* – 1}, as the fraction of pairs of polymorphic sites separated by *d* bases for which there is a mutation with derived allele count *i* at one site and a second mutation with derived allele count *j* at the other site. Note that *ϕ*_*ij*_ (*d*) = *ϕ*_*ji*_ (*d*) by symmetry. The 2-SFS has been studied for non-recombining sites by Ferretti *et al*. (2018) in a neutral model and by Xie (2011) in a model with selection. When the ancestral allele is unknown, we define the folded 2-SFS, *φ*, by analogy to the folded SFS: *φ*_*ij*_ (*d*) represents the fraction of pairs of sites separated by *d* bases in which one site has minor allele count *i* and the other has minor allele count *j* (1 ≤ {*i, j*} ≤ ⌊*n*/2⌋). For non-recombining sites, the 2-SFS is independent of the distance, so we will suppress the *d* in our notation when considering the nonrecombining case.

In the limit of low per-site mutation rate (*µ* → 0) and no recombination, all polymorphic sites are bi-allelic and the expected SFS and 2-SFS are related to moments of the genealogical branch length distribution by

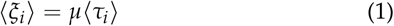

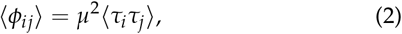

where *τ*_*i*_ is the total length of branches subtending *i* leaves of a gene genealogy and ⟨µ⟩ represents the expectation over the distribution of gene genealogies defined by a coalescent model. Thus, the SFS and 2-SFS depend on the distribution of coalescent times as well as the distribution of tree topologies. In the opposite limit of high recombination between sites (i.e., fully unlinked loci), the genealogies of the sites come from independent draws of the generating coalescent model, and the 2-SFS can be determined directly from the SFS: ⟨*ϕ*_*ij*_ ⟩ = *µ*^2^⟨*τ*_*i*_ ⟩⟨*τ*_*j*_ ⟩ = ⟨*ξ*_*i*_ ⟩⟨*ξ*_*j*_ ⟩. Thus, for a recombining population, the 2-SFS is a function of the genomic distance *d* between sites and only contains information not found in the SFS for nearby, linked sites.

Fu (1995) calculated the first and second moments of the branch-length distribution for a non-recombining infinite-sites locus under the standard time-homogeneous Kingman coalescent. He found that ⟨*τ*_*i*_*τ*_*j*_ ⟩ < ⟨*τ*_*i*_ ⟩⟨*τ*_*j*_ ⟩ for all *j* ∉ {*i*, (*n* − *i*)}. This result, combined with Eq. (1) and Eq. (2), implies a negative correlation between mutations at different frequencies: trees generating a mutation with derived allele count *i* are less likely than average to generate a second mutation with derived allele count *j* ∉ {*i*, (*n* − *i*)}. (There are positive correlations between mutations at complementary frequencies induced by genealogies whose root node partitions the tree into subtrees of size *I* and *n* − *i*.)

Birkner *et al*. (2013) extended Fu’s calculation to a family of multiple-merger coalescents called beta coalescents. This one-parameter family interpolates between the Kingman coalescent and the Bolthausen-Sznitman coalescent (Bolthausen and Sznitman 1998) as the parameter, *α*, ranges from 2 to 1. Beta coalescents arise in models with fat-tailed offspring distributions (Schweinsberg 2003; Steinrücken *et al*. 2013), and the Bolthausen-Sznitman coalescent is the limiting distribution of genealogies in populations that are rapidly adapting or experiencing extensive purifying selection (Neher and Hallatschek 2013). The calculations of Birkner *et al*. (2013) show positive correlations between *ξ*_*i*_ and *ξ*_*j*_ for *j* ∈ {*i, n* − *i*} (Figures 5 and 6 of Birkner *et al*. (2013). Thus, unlike the standard Kingman coalescent, the beta coalescent can generate positive associations between mutations with different minor allele counts. Together, these results suggest that the differences in associations between mutations at different frequencies (i.e. differences in the 2-SFS) can be used to distinguish multiple-merger coalescents from the Kingman coalescent.

### A 2-SFS-based test for the Kingman coalescent

Motivated by this reasoning, we developed a method to use information in the 2-SFS to determine whether a Kingman coalescent (with any demographic history) is consistent with realworld genomic data. The basic idea is to first use the observed SFS to determine the best-fit demographic history within the Kingman model. We then simulate the expected 2-SFS predicted by this best-fit Kingman demographic history, and use a goodness-of-fit statistic to determine whether this expected 2-SFS is consistent with the data. We illustrate this pipeline in Fig. 1 and describe each step in more detail below.

**Figure 1.**
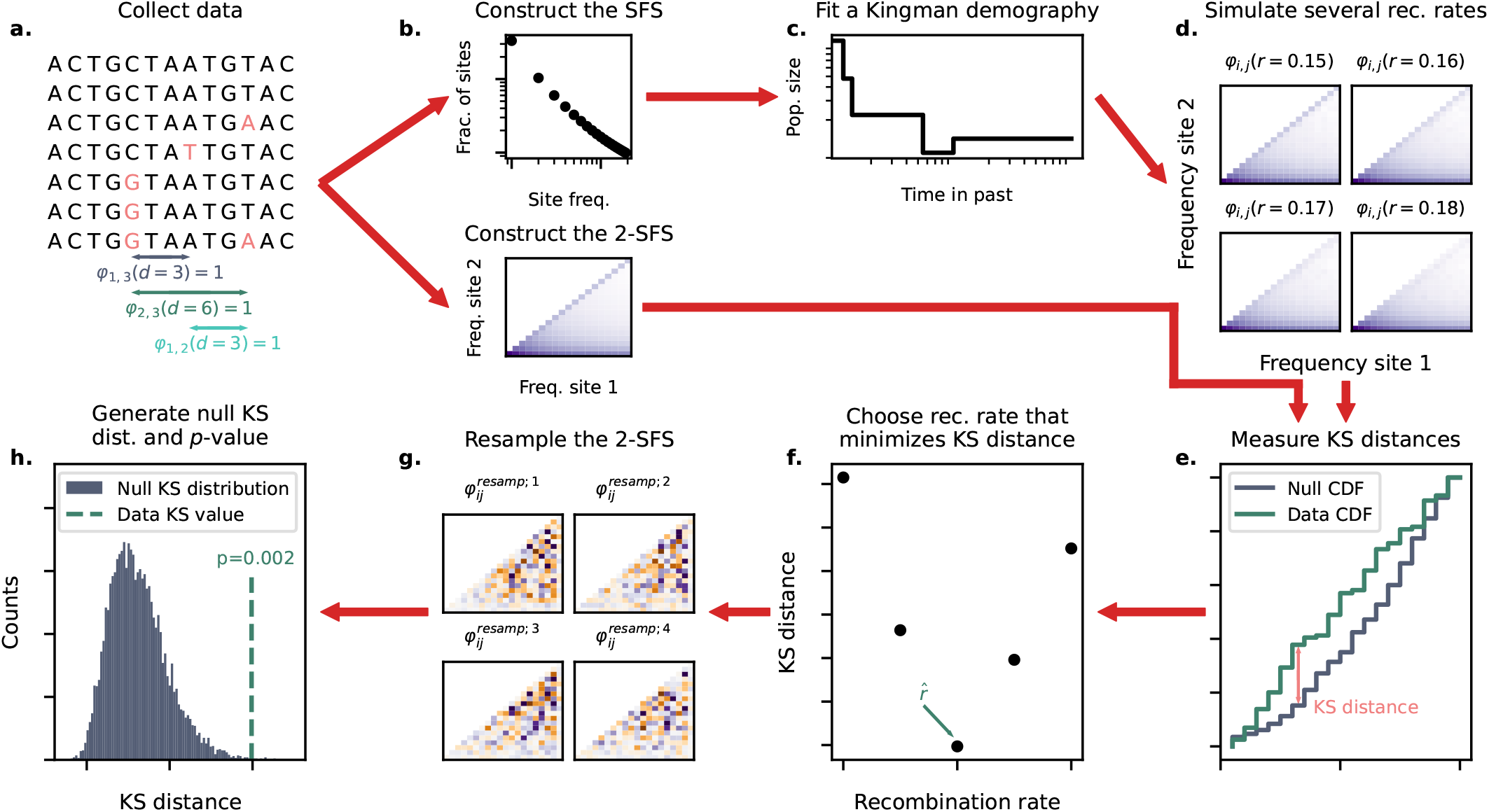
Schematic of the model-checking pipeline. The pipeline follows the red arrows from **a** to **h**. Briefly, after data collection and cleaning (**a**), we construct the SFS and 2-SFS (**b**) and fit a Kingman demography (the null model) to the SFS (**c**). We simulate the 2-SFS expected from a Kingman model with this null demography for several values of the recombination rate and choose the recombination rate, 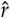, that minimizes the KS distance between the 2-SFS of the data and the null (**d-f**). We then resample 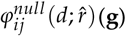 and compute the KS distsance between these resampled distributions and 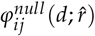 to generate a null KS distribution. We compare the KS distance between 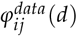 and 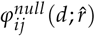 to this null KS distribution to generate a *p*-value (**h**).

#### Computing the SFS and 2-SFS from population data

We begin by generating the folded SFS and 2-SFS, 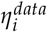 and 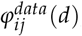, using sequence data from a sampled population. Because in practice we will often restrict our analysis to patterns at fourfold degnerate sites, we typically only consider values of *d* that are multiples of 3. If the ancestral allele is known, the unfolded SFS and 2-SFS can be used instead. To increase computational efficiency, we lump alleles with frequency larger than *k*_*max*_ into one high-frequency bin, choosing *k*_*max*_ by eye such that the high-frequency tail of the SFS is low noise (though we note that the pipeline is robust to the exact choice of *k*_*max*_, and can be implemented without this lumping if preferred).

#### Inferring the best-fit Kingman demography

We next use the observed SFS to infer the best-fit Kingman demographic model. To do so, we fit a 5-epoch piecewise-constant Kingman demography, *N*_*null*_(*t*), to the lumped SFS of the data using a modification of the `fastNeutrino` algorithm (Bhaskar *et al*. 2015). As in `fastNeutrino`, we find the *N*_*null*_(*t*) that minimizes the Kullback–Leibler (KL) divergence between the expected and observed SFS using the L-BFGS-B algorithm with automatic differentiation. Unlike `fastNeutrino`, we apply *L*_2_ regularization to the vector of log population sizes. Regularization helps the solver find well-behaved solutions by penalizing very short epochs with very large population sizes, which do not affect the SFS. Python implementation of the fitting algorithm, which we refer to as `fitsfs`, is available in a Github repository at https://github.com/desai-lab/twosfs.

Our choice of this 5-epoch model is designed to be conservative in allowing for highly flexible Kingman population histories, as compared to more restrictive assumptions such as a piecewise constant model with only one or two epochs, or models which make assumptions about the shape of past population growth. As we will see below, the inferred 5-epoch Kingman demographic model is typically an excellent fit to the observed SFS, even when the underlying model is very different (this is precisely why the SFS alone has limited power to test the Kingman assumptions).

#### Null 2-SFS and recombination rate

Once the best-fit demography has been inferred from the SFS, we generate the 2-SFS predicted by the Kingman coalescent with that demography, which we refer to as 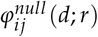, by simulating genealogies using `msprime`. We note that this predicted 2-SFS depends on the recombination rate *r*, which determines how quickly the 2-SFS decays towards the product of the corresponding SFSs as a function of *d*. However, the correct choice of recombination rate may often be unknown. To be conservative in the face of this uncertainty in the recombination rate, we therefore simulate multiple candidate null 2-SFS with different recombination rates, and choose the recomination rate that *minimizes* our ability to reject the Kingman model (as described in more detail below).

#### Statistic for comparing expected and observed 2-SFS

We next wish to compare the expected 2-SFS under the best-fit Kingman demographic model, 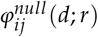, to the 2-SFS observed in the data, 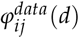. To do so, we use a form of the Kolmogorov–Smirnov (KS) distance (Kolmogorov 1933; Smirnov 1948) generalized to three variables (*i, j*, and *d*; we treat *r* as a constant here), by implementing the procedure described in Gosset (1987). The multidimensional KS distance is a nonparametric statistic that measures the degree to which an empirical distribution (here 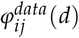) matches a proposed generating distribution (here 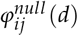). In summary, it is the maximum absolute distance between the cumulative distribution functions (CDFs) generated by 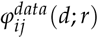 and 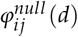, maximized again over all eight cumulation directions when defining the multidimensional CDF (i.e. Pr(*i* ≤ *I* ∧ *j* ≤ *J* ∧ *d* ≤ *D*), Pr(*i* ≥ *I* ∧ *j* ≤ *J* ∧ *d* ≤ *D*), etc). We direct readers to Gosset (1987) for a more thorough description of this statistic.

As noted above, 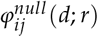 depends on the recombination rate, which is often unknown. We therefore compute this multidimensional KS statistic to compare 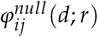 with 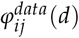 for a range of different values of *r*. We then choose the value of the recombination rate, 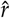, which minimizes this KS distance between the 2-SFS expected under the Kingman demographic model and the 2-SFS observed in the data. This ensures that we are conservative in rejecting the Kingman model in the face of uncertainty about the recombination rate.

To efficiently find 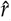, we choose candidate recombination rates using a golden-section search (Kiefer 1953). Starting with conservative lower and upper bounds for 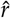, the golden-section search algorithm iteratively proposes new candidate recombination rates and narrows the bounds on 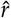 through sequential evaluations of the KS distance between 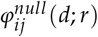 and 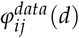. The algorithm can be run for a given number of iterations or until some other stopping criteria is met; in this paper, we run the algorithm for five iterations.

#### Null KS distribution and p-value

We next wish to determine whether the observed multidimensional KS distance is consistent with 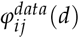 being drawn from 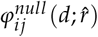. In other words, is the observed 2-SFS consistent with the 2-SFS expected based on the best-fit Kingman demographic model? The complex natures of our KS statistic and the noise associated with mutation accumulation and population sampling mean that it is not possible to derive an analytic expression for a range of “typical” KS distances to be expected assuming the null model is correct. Therefore, we approximate the null KS distribution through a resampling procedure. Specifically, we generate 1000 candidate 2-SFS distributions, 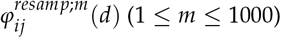, by drawing *PD*(*d*) multinomial samples at each genomic distance *d* 1000 times from 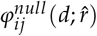, where the pair density *PD*(*d*) is the number of pairs of sites at distance *d* in the sample. That is, every iteration of 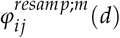 has the same number of pairs of sites at distance *d* as the sampled data, but with expectation value equal to 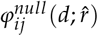. Intuitively, these resampled 2-SFS distributions can be thought of as versions of the null 2-SFS “noised” to the level of the sampled data. By calculating the KS distance between each 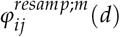 and 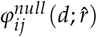, we generate an approximate null KS distribution to which the KS distance between 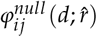 and 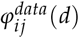 can be compared. We then use this comparison to generate a *p*-value for the rejection of the Kingman model.

#### Model-checking pipeline

We have implemented this 2-SFS based model checking procedure in a `Snakemake` pipeline which can be used to test whether any real-world or simulated population data is consistent with the Kingman coalescent, publicly available in a Github repository at https://github.com/desai-lab/twosfs. This repository has code to reproduce all results and figures from this manuscript and is straightforward to edit to test parameter values ouside those explored in this paper. Users wishing to test real-world data using the pipeline must supply a `JSON` file containing the locations of all polymorphic sites and their associated derived or minor allele counts. This file uses a specific custom format, though we supply code for conversion from both `VCF` and text file formats. The pipeline further requires an upper and lower bound for the recombination rates and contains flags for various data cleanup choices. We direct readers to the `README` located in our Github repository for further details and instructions.

### Validation of our 2-SFS based test with simulations

To test the performance of our model checking procedure, we simulated coalescent histories using `msprime` (Baumdicker *et al*. 2022) and `SLiM` (Haller *et al*. 2019) under four classes of models: (1) the neutral, constant-size Kingman coalescent; (2) a neutral, exponentially growing Kingman coalescent; (3) a neutral, constant-size beta coalescent; and (4) a constant-size population undergoing positive selection at many sites along the genome. For each type of simulation, we tested a range of relevant parameter values, as shown in Table 1.

**Table 1.** Models and parameter ranges for simulated coalescent processes.

| Model | Parameter range |
| --- | --- |
| Constant-size,<br>neutral Kingman | N/A |
| Exponential<br>growth | Growth rate $\gamma$: 0.25 – 2.0 per $T_2$<br>Growth time $t_0$: 0.5 – 2.0 * $T_2$ |
| Beta coalescent | $\alpha$: 1.05 – 1.95 |
| Positive Selection | Selective coefficient $s$: 0.005 – 0.08%<br>Rate of selective mutations $\mu$: $10^{-10}$ – $10^{-11}$ per site per genome per genera-<br>tion<br>Diploid population size $N = 10,000$<br>Genome length $L = 5,000$ |

#### A flexible Kingman demography reproduces features of a non-Kingman SFS

For every model-parameter combination, we first simulated the expected folded SFS of 100 samples. Because of the stochasticity at higher frequencies, we combined all mutations with frequency *k* ≥ *k*_*max*_ = 20 into one “lumped” high-frequency bin. As described above, we then used `fitsfs` to fit a piecewise-constant neutral demographic model to each lumped, folded SFS. We show one example of the resulting SFS from each of the four types of models we simulated, along with the corresponding `fitsfs` fits, in Fig. 2. We see that the observed site frequency spectra deviate strongly from the constant-size Kingman expectation for the three examples where this was not the underlying model. However, we find that a Kingman coalescent with a flexible population size can be fit to all four spectra nearly perfectly. This implies that any statistics based solely on the SFS, or transformations thereof, will have minimal power to distinguish the non-Kingman scenarios (here beta coalescent and positive selection models) from a sufficiently flexible Kingman demography.

**Figure 2.**
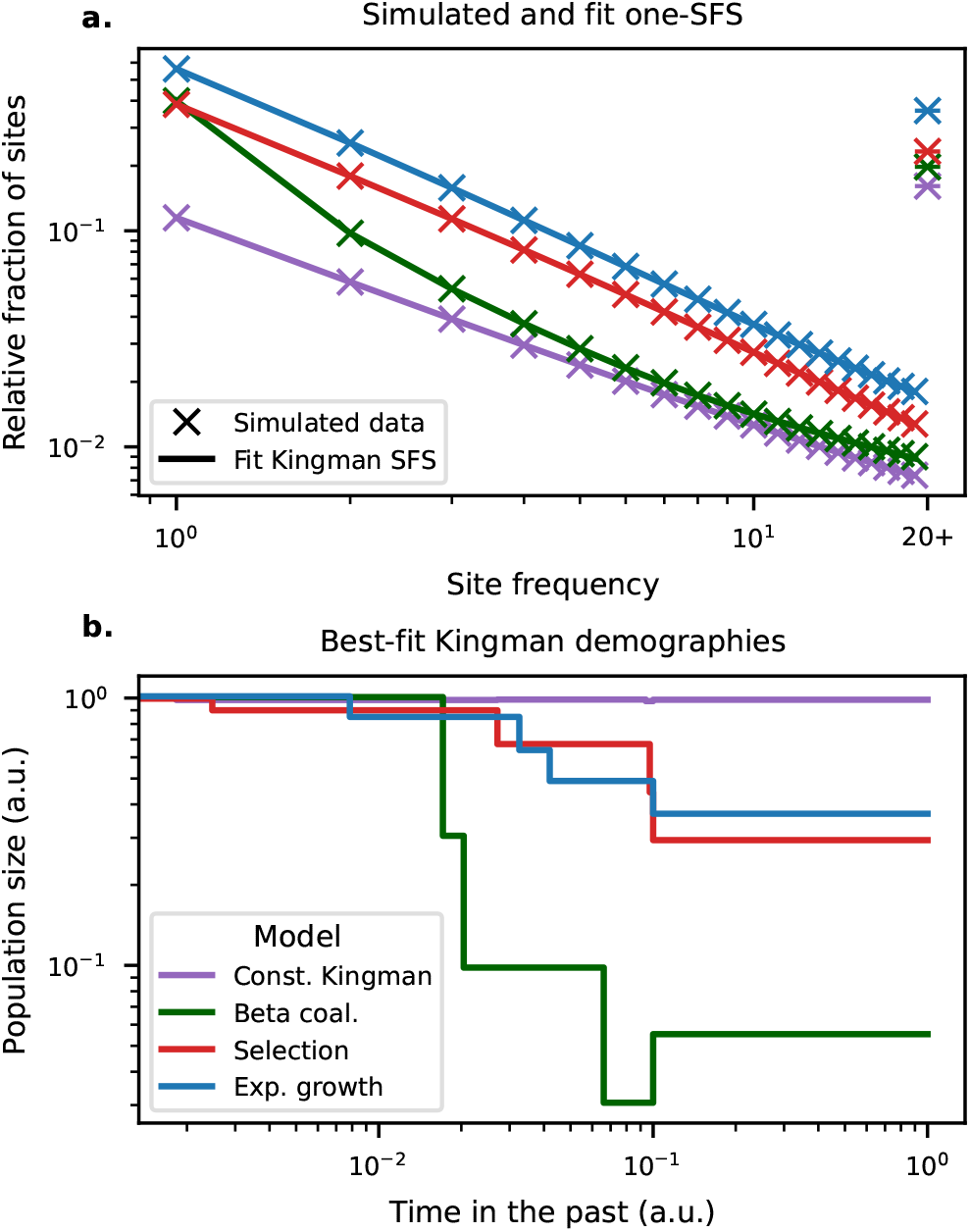
(**a**) Simulated site frequency spectra for example constant-size-Kingman, beta coalescent, positive-selection, and exponential population size growth models, compared to the expectations from the corresponding best-fit Kingman demographic models. The examples shown here are for *α* = 1.3 (beta coalescent); *s* = 0.55, *µ* = 10^−10^ (positive selection); and *t*_0_ = 0.5, *γ* = 2.0 (exponential growth). Note that site frequency spectra are shifted vertically relative to each other to aid in visibility. (**b**) The inferred best-fit Kingman demographic models for each of the four examples shown in (**a**). Population size and time in the past have units of an arbitrary coalescent timescale.

#### The 2-SFS can distinguish demographic models with matching SFS

By contrast, we expect that the 2-SFS should allow us to distinguish non-Kingman scenarios from a Kingman demographic model that generates an identical SFS. To show this, we used `msprime` to simulate the Kingman coalescent with the piecewise-constant demographic histories inferred by `fitsfs` for the simulated models described above. This produced a set of pairs of simulations, each consisting of an original (potentially non-Kingman) model, along with the corresponding Kingman model with the piecewise-constant demographic history that is the best fit to the SFS from the original model.

By construction, these simulated Kingman coalescents produce nearly identical SFS as the corresponding original models. We then compared the 2-SFS produced by these simulated Kingman coalescents to those produced by our simulations of the original models. To visualize this comparison, in Fig. 3 we plot four examples of the log-ratio of the 2-SFS produced by the original models to those produced by the best-fit Kingman demographic model. We see that for the beta and positive selection cases, where the original model is not Kingman, there is a striking visual difference with the 2-SFS of the corresponding Kingman demographic model, despite the near-perfect fit to the SFS. On the other hand, the constant-size and exponentially growing Kingman coalescents show signal consistent with simulation noise. Taken together, these results imply that the 2-SFS can distinguish between Kingman and non-Kingman coalescent models, even when the SFS fails to do so.

**Figure 3.**
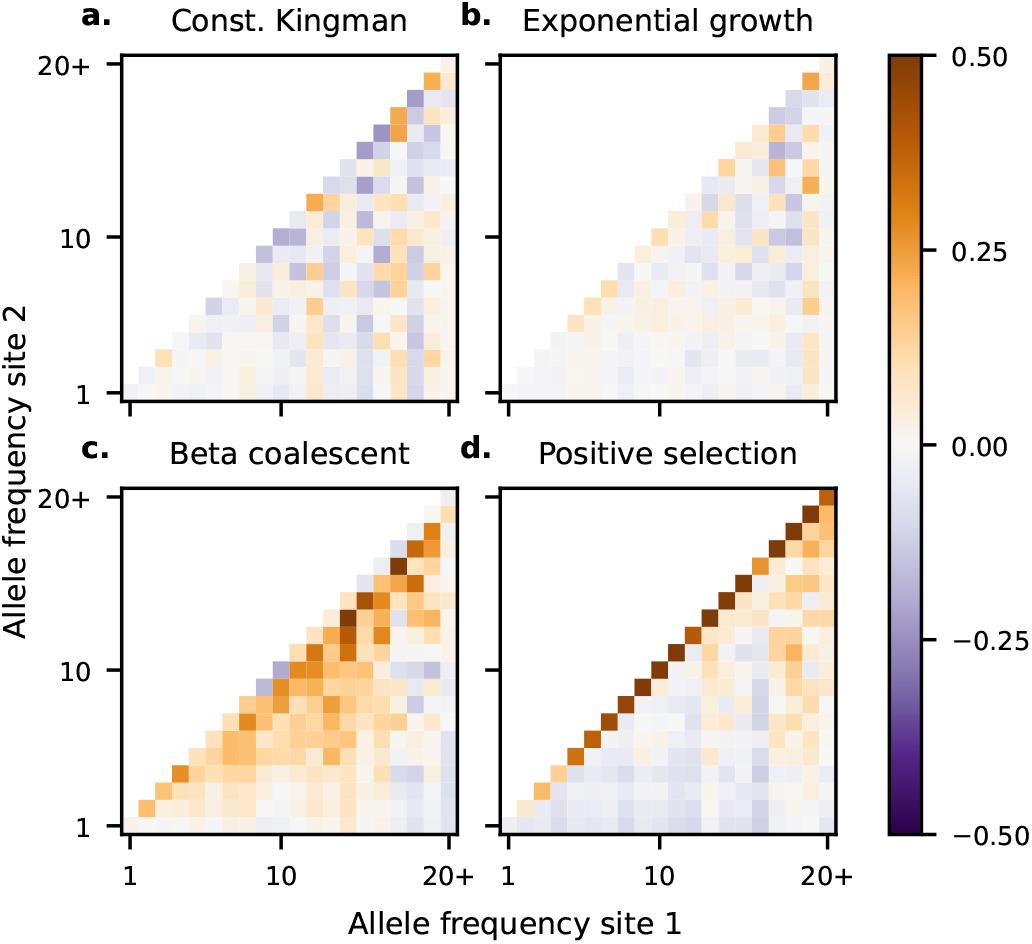
Log-ratio of the 2-SFS of the four example models shown in Fig. 2 with the 2-SFS expected under the corresponding best-fit piecewise-constant Kingman demographies.

#### Power analysis of our model checking procedure

The examples shown above demonstrate visually that there is information in the 2-SFS that can potentially be used to distinguish Kingman from non-Kingman coalescent processes. To determine whether the statistical test we introduced above effectively uses this information, we validated our model-checking pipeline with the simulated models from Table 1. Proper validation requires several replicate simulated SFS and 2-SFS (i.e. multiple simulations of *η*^*data*^ and *φ*^*data*^), which are computationally expensive to generate from scratch for the large number of models we simulate. Therefore, to save computational resources, we re-employed the resampling method described earlier. For each model-parameter combination, we generated 100 simulated 2-SFS, *φ*^*sim*;*l*^, by resampling the low-noise 2-SFS 100 times at *PD*(*d*) = 10, 000 for genomic distances *d* = {3, 6, …, 24}, approximately matching the pair density of fourfold degenerate sites in the *D. melanogaster* dataset we describe below. Again, each of these *φ*^*sim*;*l*^ can be thought of as a 2-SFS whose expectation value matches the simulated coalescent model but is noised to mimic real-world data. We ran our model-checking pipeline independently for each *φ*^*sim*;*l*^, generating 100 validation runs of the procedure for every model-parameter combination.

We plot the power to reject Kingman coalescence at a *p*-value threshold of 0.05 in Fig. 4. As seen in Fig. 4**a-c**, we have high power to reject Kingman coalescence for models that in-volve highly skewed offspring distributions and strong positive selection and low false-rejection rates for neutral exponential growth for biologically realistic sample sizes. In other words, we correctly reject Kingman coalescence whenever the underlying model involves sufficiently strong non-Kingman processes, but do not incorrectly reject the model in any of the scenarios involving exponential growth. This trend holds despite the three model classes spanning similar levels of distortion of the SFS, as measured by Tajima’s *D* (Fig. 4**d**).

**Figure 4.**
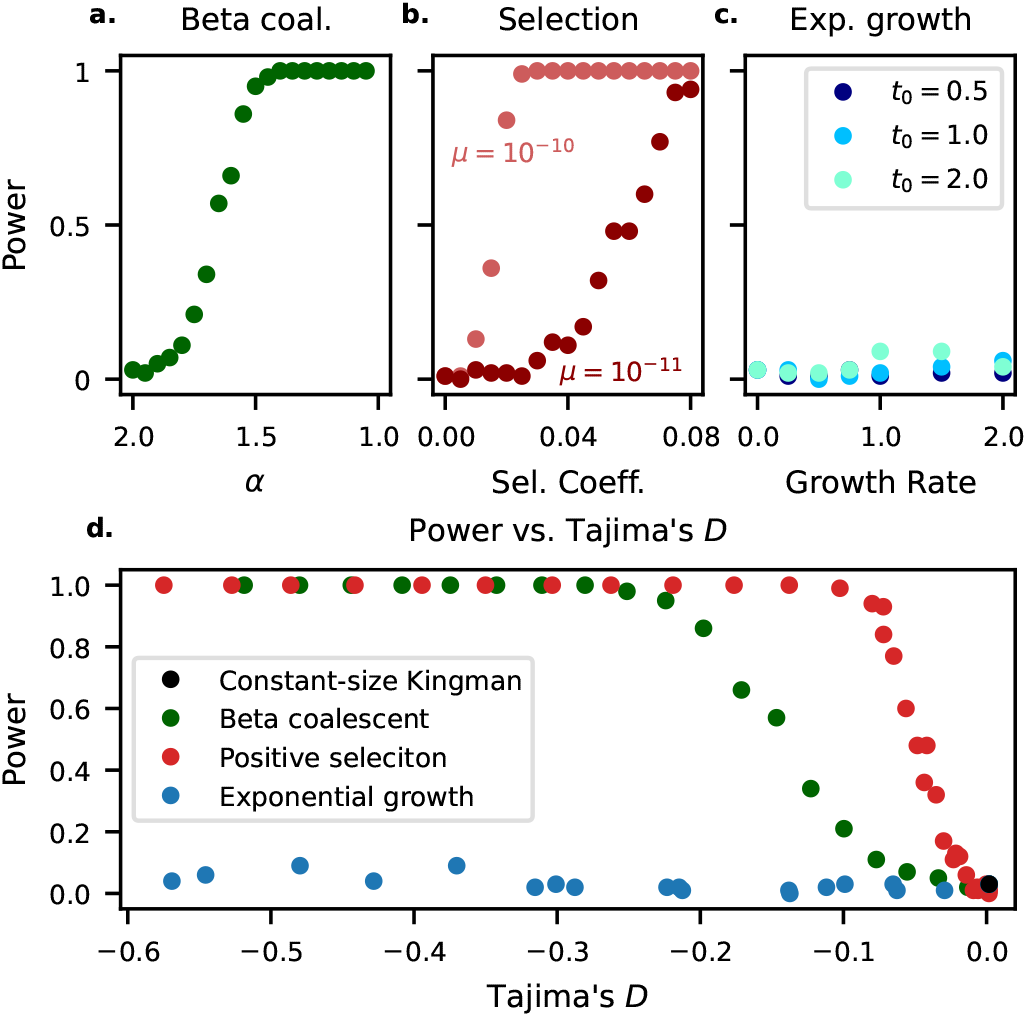
(**a-c**) Power to reject Kingman coalescence in simulations across a range of parameter values for several different classes of models. For the beta coalescent and positive selection, power increases as simulations move away from neutrality, as expected. In exponentially growing Kingman coalescents, false rejection rates remain low for all parameter values. (**d**) Tajima’s *D*, which is a measure of the degree to which the SFS is distorted relative to its expectation under a constant-size Kingman model, versus power to reject Kingman coalescence. Note that our model-checking pipeline demonstrates high power to detect non-Kingman coalescence and low false-rejection rates for Kingman models with non-constant population size history, despite similar distortions to the SFS as measured by Tajima’s *D*.

### Analysis of *D. melanogaster* data

We next applied our method to analyze sequence data from the DPGP3 data set, which consists of haploid consensus sequences from ∼150 flies, obtained via the haploid embryo method of Langley *et al*. (2011). The SNP calls that characterize these sequences were subjected to a variety of quality filters as described in Lack *et al*. (2015). We obtained the DPGP3 consensus sequence files version 1.1 for the 2L, 2R, 3L, and 3R chromosome arms from www.johnpool.net/genomes.html. These files contain sequence alignments of all flies in the sample on all chromosome arms. We also downloaded the Nov. 3, 2016 spreadsheet of inversions available at the same link. For each chromosome arm, we excluded any samples with an inversion in that arm and then randomly down-sampled to *n* = 100 flies. As a result, the data for each chromosome arm is from a different subset of individuals.

To ensure our analyses focused on putatively neutral variation, we filtered called SNPs to fourfold degenerate sites. We then calculated the average pairwise diversity, Π, as a function of position for each autosomal chromosome arm. Pairwise diversity is high in the middle of each chromosome arm and lower near the centromeres and telomeres, in agreement with calculations by Corbett-Detig *et al*. (2015). Our modeling – and coalescent-based demographic inference in general – assumes that the distribution of gene genealogies is homogeneous along the chromosome. Therefore, we selected a 13-16 Mb “central” region of each arm with relatively homogeneous values of Π for further analysis. The boundary positions of these central regions are given in Table 2.

**Table 2.** Central regions and fraction of sites above 90% coverage for the four *D. melanogaster* chromosome arms analyzed in this study. The cutoff positions of central regions are referenced to the DPGP3 reference genome available at http://www.johnpool.net/genomes.html.

| Chromosome arm | Central region analyzed | Fraction of sites kept |
| --- | --- | --- |
| 2L | 1 – 17 Mb | 0.916 |
| 2R | 6 – 19 Mb | 0.928 |
| 3L | 1 – 17 Mb | 0.928 |
| 3R | 10 – 26 Mb | 0.912 |

In order to ensure that the segregating mutations reflect true genetic diversity and not variation in calling errors, we excluded sites with fewer than 90 of the 100 genotypes called. This leaves over 90% of all sites and does not substantially alter the fraction of polymorphic sites (Table 2). For remaining sites with missing calls, we probabilistically imputed missing genotypes as either the major or minor allele based on the proportion of called genotypes at that site. Every missing read was assigned the minor allele with probability *p* and the major allele with probability 1 − *p*, with *p* equal to the minor allele fraction of called genotypes at that site.

We ran each chromosome arm independently through our model testing pipeline, fitting Kingman demographies to the SFS and comparing the 2-SFS of the data to the fit demographies. We plot these results in Fig. 5. For all chromosome arms, the best-fit Kingman demographies closely match the observed SFS (Fig. 5**a**) and show a recent roughly doubling of the population size (Fig. 5**b**). However, the 2-SFS of our inferred demographies do not match the 2-SFS observed in the data (Fig. 6), as can be seen visually (Fig. 6**a-d**) and verified numerically using our KS statistic (Fig. 6**e**). This implies that this *D. melanogaster* data is inconsistent with the Kingman coalescent, and that the best-fit Kingman demographies are not an accurate representation of the effective population size history but are instead fitting the effects of other types of non-Kingman processes.

**Figure 5.**
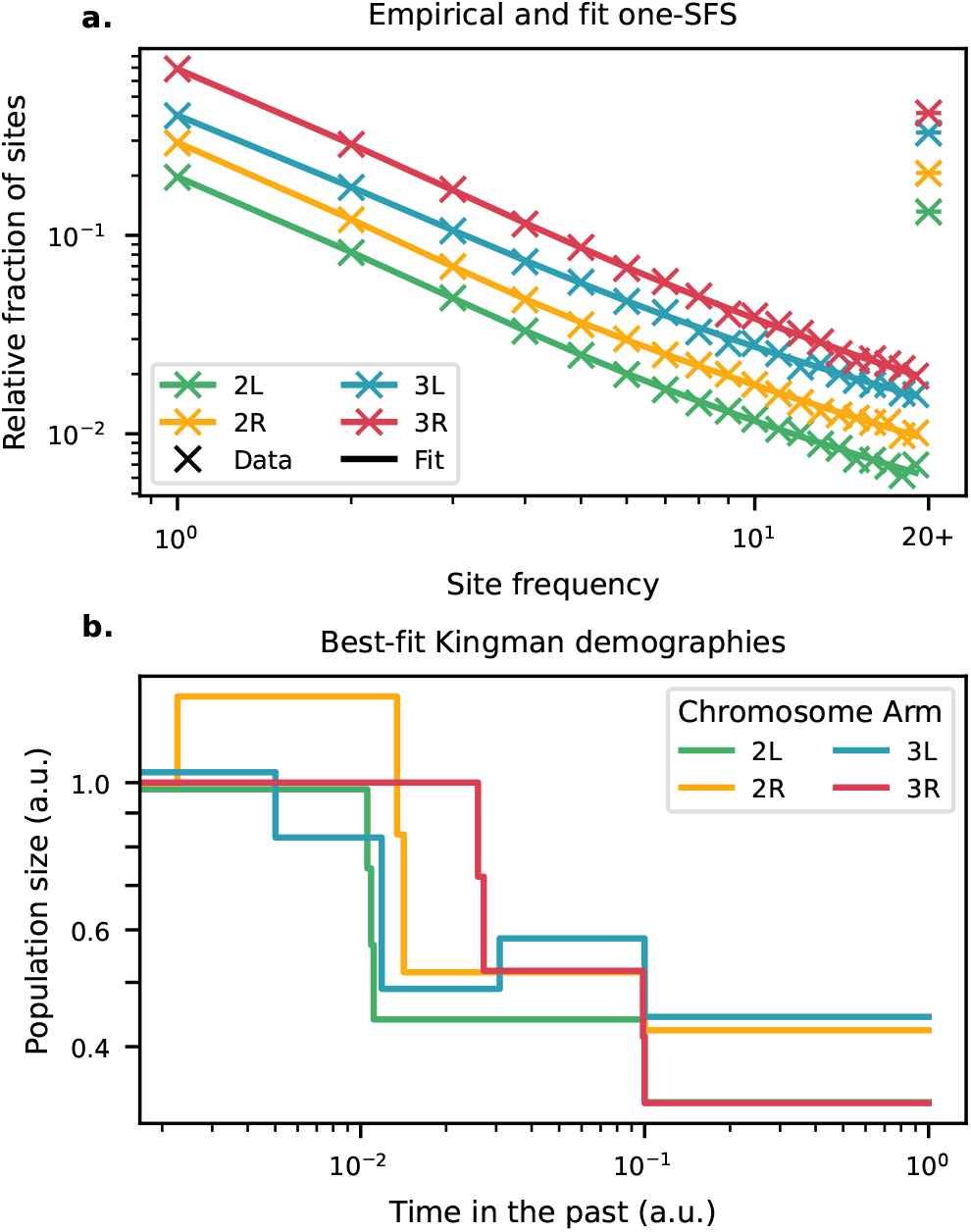
(**a**) Observed and fit SFS for the four *D. melanogaster* chromosome arms investigated in this study. Note that SFS of each chromosome arm are shifted vertically to improve visibility. The SFS of the fit demographies closely match those from the data. The demographic models that produce the fit site frequency spectra are plotted in (**b**).

**Figure 6.**
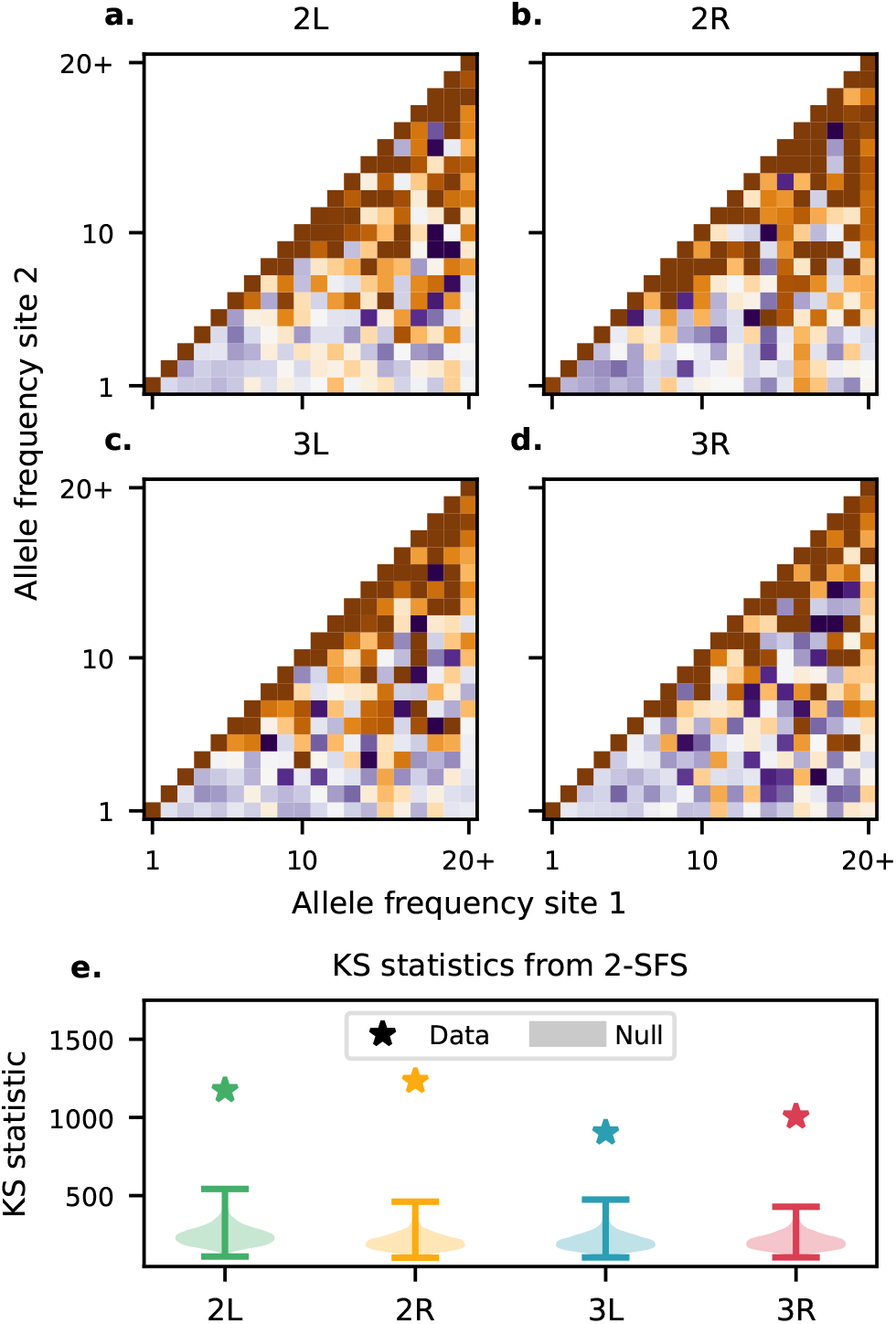
(**a-d**) Log-ratios between the observed 2-SFS and the 2-SFS expected from a Kingman coalescent fit to the SFS for each of the *D. melanogaster* chromosome arms investigated in this study. Note the clear visual mismatch between the observed and expected 2-SFS. (**e**) Empirical null KS distributions (shaded regions) and measured KS distances between the data and the Kingman fit (stars) for each of the chromosome arms investigated. All 2-SFS deviate significantly from the null distributions (*p* < 10^−3^).

## Discussion

We have shown that the 2-SFS is sensitive to multiple mergers, but largely invariant to population growth in the Kingman coalescent, making it well-suited for coalescent model checking. We developed and validated a model-checking procedure that uses this information to discriminate Kingman from non-Kingman coalescence, and demonstrated the power of our approach in simulated data. We then applied this method to data from *D. melanogaster*, which is believed to be strongly shaped by natural selection, and found evidence that population growth alone cannot explain the correlation structure in the 2-SFS in this system.

We emphasize that our 2-SFS-based test is fundamentally different from approaches based on the SFS or on statistics derived from the SFS. For example, several recent studies have developed methods to use the SFS to distinguish multiple-merger coalescents from Kingman models with specific forms of population growth (Birkner *et al*. 2013; Blath *et al*. 2016; Spence *et al*. 2016; Eldon *et al*. 2015; Koskela 2018; Koskela and Wilke Berenguer 2019). In contrast to this work, our approach uses the SFS to infer the best-fit Kingman demographic model, and then asks whether this best-fit Kingman model is consistent with the different information contained in the *joint* frequency spectrum of pairs of sites. The relationship between our method and other more recent work (Freund and Siri-Jégousse 2021; Menardo *et al*. 2021; Johri *et al*. 2020; Lepers *et al*. 2021) that uses an approximate Bayesian computation framework to distinguish between coalescent models is more complex. These studies make use of several SFS-derived statistics as well as additional statistics related to clade size and linkage disequilibrium. These additional statistics are not directly related to the 2-SFS but may contain some related information.

We can get an intuitive understanding for why 2-SFS-based statistics are useful in distinguishing between coalescent models by considering how the 2-SFS depends on the distribution of branch lengths and tree topologies. Mathematically, the expected 2-SFS, 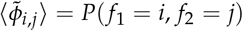 can be directly related to the set Ψ of tree topologies allowed by the coalescent model:

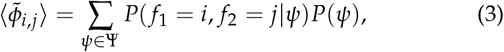

where *f*_{1,2}_ denote the frequencies of mutations at some sites 1 and 2 and *ψ* is a particular tree topology. We note here that the first term in this equation, *P*(*f*_1_ = *i, f*_2_ = *j*|*ψ*), depends only on the distribution of branch lengths (which can be manipulated arbitrarily using an appropriate choice of historical population size). On the other hand, the second term, *P*(*ψ*), reflects only the distribution of tree topologies, which depends heavily on the particular coalescent model. This expression for the 2-SFS can be further expanded as:

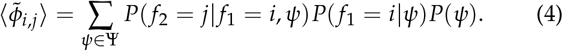

Using Bayes’ Theorem, we can rewrite this as:

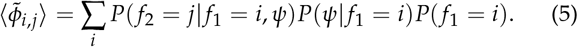

We note again that the first term, *P*(*f*_2_ = *j*|*ψ, f*_1_ = *i*), depends only on the distribution of branch lengths, while the last term, *P*(*f*_1_ = *i*), is just the expected SFS, ⟨*ξ*_*i*_ ⟩. As argued above, by allowing the population size (and thus the coalescent rate) to be explicitly time-dependent, the SFS can be made arbitrarily similar between the Kingman coalescent and broad classes of multiple-merger coalescents. Therefore, we find a condition for two coalescent models to be theoretically distinguishable using the 2-SFS:

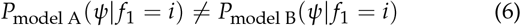

In summary, the dependence of the 2-SFS on tree topologies contains a term that depends on the coalescent model but not on the SFS. In other words, the 2-SFS distinguishes between models with identical SFS when the trees used to generate the SFS differ between models.

We can further see from the above discussion why the 2-SFS is particularly useful in distinguishing Kingman from multiple-merger coalescents. In any coalescent model, the presence of a site at frequency *i* implies that there is a branch in the coalescent history that subtends *i* leaves. However, given this, the probability that the next coalescent event creates a branch that subtends *k* of these *i* leaves is uniformly distributed in the Kingman model, while more skewed offspring distributions can be created by multiple merger events. These types of effects mean that the probability of a given topology conditional on observing the mutation at frequency *i* can differ substantially between Kingman and multiple-merger models.

Throughout this study, we have focused on developing a statistical test that allows us to reject Kingman coalescents with flexible time-dependent population size histories. We have analyzed the power of this method when the true population history involves either a beta coalescent or recurrent positive selection. However, these are far from the only genealogical models that may describe a population’s history, and researchers may be interested in discriminating arbitrarily between these models, rather than simply rejecting a Kingman coalescent. For example, the differences in the 2-SFS produced by the beta coalescent and positive selection (Fig. 3) suggest that it may be possible to use 2-SFS based statistics to discriminate between these two models. More generally, extending our framework to allow for comparison between two or more arbitrary coalescent models is an exciting area for future work.

We have also focused in this study on a single application of our statistical test to data from *Drosophila melanogaster*. However, there are a broad range of possible further empirical applications. For example, one interesting direction would be to use 2-SFS-based statistics to assess the evidence for variation in multiple-merger coalescence within genomes and between species, potentially identifying genomic regions and organisms that are more likely to be under strong selection. Alternatively, one could survey multiple species using a data set such as the diversity data compiled by Corbett-Detig *et al*. (2015). These are interesting avenues for future work, which hold the potential to reveal new information about the suitability of widely used population genetic models, and could provide further insight into the forces that determine genetic diversity.

## Acknowledgements

We thank Arjun Biddanda, Maryn Carlson, Ivana Cvijović, Ben Good, Dick Hudson, Evan Koch, Joe Marcus, Richard Neher, Matthias Steinrücken, John Wakeley, and Aleksandra Walczak for helpful discussions and comments on the manuscript. DPR was supported by the Chicago Fellows Program of the University of Chicago. JN acknowledges support for this work from NIH grants GM108805 and HG007089. MMD acknowledges support from grant PHY-1914916 from the NSF and grant GM104239 from the NIH. This work was completed in part with resources provided by the University of Chicago Research Computing Center and the Harvard Faculty of Arts and Sciences Research Computing Center.

